# Pharmacological modulation of ATP-binding cassette transporters alters feeding phases of the sugarcane aphid on susceptible sorghum

**DOI:** 10.64898/2026.09.05.749624

**Authors:** Leslie C. Rault, Kashish Verma, Lise Pingault, Joe Louis, Troy D. Anderson

**Author notes:** Sharing first authorship (Leslie C. Rault, Kashish Verma).

## Abstract

**BACKGROUND:** Aphids damage crops through sustained feeding and by rapid transmission of plant viruses, processes that are not effectively prevented by conventional insecticides. The sugarcane aphid (*Melanaphis sacchari*), an invasive pest of sorghum in the United States, can inoculate non-persistent viruses within minutes of probing. Disrupting feeding phases represents an alternative strategy in plant disease and integrated pest management. ATP-binding cassette transporters (ABC transporters) influence insect physiological performance, but their role in regulating aphid feeding behavior relevant to crop damage and virus transmission remains unknown. This study evaluated whether ABC transporter activity alters feeding phases of *M. sacchari* on sorghum.

**RESULTS:** Aphids exposed to ABC transporter modulators via artificial diet and monitored using Electrical Penetration Graph (EPG) recordings showed significant reductions in salivation in E1 phase, compared with controls (P < 0.05) and an increased duration in xylem phase when treated with elacridar, an ABC transporter inhibitor. Aphids treated with dexamethasone, an ABC transporter activator, exhibited increased time spent in pathway phase compared to the controls. ATPase assays also showed significant variations in modulator-treated samples compared to the controls.

**CONCLUSION:** These results demonstrate that transporter modulators alter feeding behaviors which might lead to reduced crop damage/virus transmission, including pathway activities, salivation, and xylem ingestion. This study opens the path to identify ABC transporters as regulators of feeding phases, and to advance the understanding of aphid-plant interactions. This new knowledge will support the development of feeding-based pest management strategies to reduce crop damage and virus transmission.

## Introduction

Sorghum (*Sorghum bicolor* (L.) Moench) is one of the world’s most important crops, grown for grain and bioenergy. It provides food and fiber due to its high content of nutrients and phenolic compounds, as well as health-promoting properties (Dahlberg, 2019; Khalid et al., 2022). However, sorghum is susceptible to insect pests that can dramatically reduce yields via direct feeding and virus transmission (Singh et al., 2004; Vasquez et al., 2025). Aphids are particularly efficient vectors of plant viruses, with virions present on the cuticle lining of the stylet and salivary canal, allowing rapid transmission during probing and feeding on plant tissues (Brault et al., 2010). Feeding by sugarcane aphid (SCA; *Melanaphis sacchari* (Zehntner)) on sorghum can severely impair photosynthesis by removing sap from leaves and transmitting viruses, including sugarcane yellow leaf, and sugarcane mosaic viruses in sugarcane and sorghum. The combination of feeding damage and viral infection causes substantial yield losses (Bhargava et al., 1971; Bowling et al., 2016; Long and Hensley, 1972; Schenck and Lehrer, 2000; Singh et al., 2004). SCA has rapidly expanded its geographic and host range in the United States, infesting 23 states (EDDMapS, 2026), where it colonizes sorghum, millet, and Johnsongrass (Bowling et al., 2016; Vasquez et al., 2025; Wright et al., 2016). Its ability to reproduce quickly and adapt to multiple hosts makes it a highly threatening pest.

Insects possess ATP-binding cassette transporters (ABC transporters), a superfamily of transmembrane proteins that transport lipophilic substrates and ions via ATP hydrolysis (Tarr et al., 2009). ABC transporters are involved in a range of biological processes, including functioning in the blood-brain barrier, acting as gut receptors for Cry toxins, mediating stress responses, and contributing to insecticide resistance (Dermauw and Van Leeuwen, 2014; Enders et al., 2020; Heckel et al., 2007). They play key roles in secondary compound detoxification in soybean aphids, *Aphis glycines* (Matsumura), feeding on aphid-resistant plants, alongside esterases and cytochrome P450s, while they are essential for development in the corn planthopper *Peregrinus maidis* (Ashmead) (Bansal et al., 2014; Wang et al., 2023).

Transcriptome studies of the Western tarnished plant bug, *Lygus hesperus* (Knight) reveal extensive ABC transporters representation (Hull et al., 2014) and ABCB6 upregulation in cotton bollworm, *Helicoverpa armigera* (Hübner) confers tolerance to gossypol (Jin et al., 2020). ABC transporters are also present in the Chinese cicada *Subpsaltria yangi* (Chen), in salivary glands, where they translocate amino acids, sugars, lipids, ions, metals, and alkaloids, facilitating adaptation to host plants (Liu et al., 2019).

These studies show the importance of ABC transporters in hemipteran feeding physiology. It is currently unknown if aphid ABC transporters in the gut or salivary glands regulate transport of sugars, amino acids, secondary metabolites, or salivary components. These functions could influence probing efficiency, phloem ingestion, and virus transmission. Similar roles have been observed in other insects, where transporter activity mediates adaptation to host chemistry and environmental stress (Liu et al., 2019; Wang et al., 2023). This mechanistic perspective supports the hypothesis that ABC transporters are integral not only for xenobiotic detoxification but also for feeding behavior and aphid performance on crops. It is not yet known how ABC transporters influence aphid feeding behavior. Our study will examine the contribution of ABC transporters to SCA feeding behavior and performance using ABC transporter modulators, providing insight into their importance for aphid feeding. Similar experiments have been conducted using potassium channel inhibitors in *Aphis gossypii* (Li et al., 2019; O’Hara et al., 2023), showing the feasibility of feeding modulation with chemical compound exposure. Understanding ABC transporter-mediated feeding mechanisms could inform the development of novel pest management strategies that reduce aphid damage and virus transmission in sorghum while minimizing non-target impacts. Strategies targeting feeding regulation rather than direct lethality may also reduce selection pressure associated with conventional insecticides.

## Materials and methods

### Plants

For experiments, BCK60 SCA-susceptible sorghum plants, *Sorghum bicolor* (L.) Moench, were grown in pots filled with soil mixed with vermiculite and perlite (PRO-MIX BX BIOFUNGICIDE + MYCORRHIZAE, Premier Tech Horticulture Ltd., Canada) at the University of Nebraska-Lincoln greenhouse with a 16-h-light/8-h-dark photoperiod, 160 µE m^-2^s^-1^, 25 °C, and 50–60% relative humidity. Plants were watered regularly and fertilized once a week. Two-week-old plants (Vanderlip and Reeves 1972) were used for all the experiments.

### Insects

A sugarcane aphid (SCA) colony was originally founded from a single wingless aphid collected from sorghum fields at the Louisiana State Agricultural Center Dean Lee Research Station, Alexandria, LA, in July 2014. A single parthenogenic female from the above colony was reared on susceptible sorghum genotype, BCK60 in a growth chamber (Thermo Scientific), at 16:8□h light: dark cycle and 25 °C at the University of Nebraska-Lincoln Department of Entomology. BCK60 plants were grown in the greenhouse to panicle initiation growth stage (Vanderlip and Reeves, 1972) and replaced degenerated plants in the growth chamber.

### Pretreatments with ABC transporter modulators in sucrose diet

Apterous adult SCA from the maintained colony were collected, starved briefly, and exposed to a 20% sucrose diet supplemented with either control solution (0.2% DMSO), elacridar (ABC transporter inactivator; TargetMol Chemicals Inc., Wellesley Hills, MA), or dexamethasone (ABC transporter activator; TCI Chemicals, Tokyo, Japan) at 0.5 mM prior to ATPase assay and EPG experiments. This concentration was chosen after an initial preliminary range finding experiment that eliminated higher concentrations that proved distasteful for SCA. Elacridar and dexamethasone are known modulators of ABC transporter activity, including ABCB, ABCC, and ABCG transporter families (Gampa et al., 2020). Aphids were starved for 3 h before being transferred to feeding chambers consisting of stretched Parafilm enclosing 700 µL of diet solution. Diets were supplemented with 2 mg/mL red dye to confirm feeding (Wille and Hartman, 2008). Only red-colored aphids that had fed successfully on the diets were used for subsequent experiments (Supplemental Figure S1).

### ATPase assay

ATPase activity was assessed as an indicator of ABC transporter function, since it relies on the energy release during ATP hydrolysis (ATPase activity). After 24 h on diet with 0.5 mM of ABC transporter modulator (the inhibitor elacridar or the activator dexamethasone) in sucrose or a control (sucrose diet), ATPase activity was measured using a commercial assay QuantiChrom™ ATPase assay kit (BioAssay Systems, Hayward, CA) to confirm compound impact on target activity, with four replicates of 10 SCA per condition tested (control and treatments). Briefly, each replicate of 10 SCA was crushed in a microcentrifuge tube with a pestle, 250 µL assay buffer was added, and the samples were centrifuged at 4º C for 5 min at 10,000 x *g*, subsequent steps followed the manufacturer’s protocol, and each reaction included 10 µL of 4 mM ATP. Absorbance was detected with a microplate reader at 620 nm and each sample measurement was normalized based on protein content, measured from an adapted protocol from Smith et al., (1985). Treatment effects were evaluated using a one-way ANOVA and a Tukey’s multiple comparison between the treatments (*P* < 0.05) using GraphPad Prism version 10.

### Monitoring of SCA feeding behavior with Electrical Penetration Graph (EPG) after exposure to inhibitors

EPG measurements were carried out in an electrically grounded Faraday cage to shield the setup from external electrical noise, at room temperature. A gold wire was attached to the dorsum of the aphid with conductive silver paint. The gold wire was connected to the insect electrode. A copper wire (plant electrode) from the EPG monitor was inserted into the damp soil of the pot.

Both electrodes were connected to a GIGA-8 EPG model (EPG Systems, Wageningen, The Netherlands) with a 10^9^ Ω resistance amplifier and adjustable voltage. Recordings of individual aphids on sorghum plants lasted 8 h and were carried out under constant light. SCA-susceptible sorghum plants, BCK60, were grown for two weeks in preparation of the EPG assay. At 24 h prior to the EPG, SCA were starved for 3 h before being placed on sucrose diet, elacridar-supplemented sucrose diet or dexamethasone-supplemented sucrose diet, as described above. Fed aphids appeared red-colored and were then starved for 1-1.5 h before introducing to the plants to promote plant feeding. Aphids were placed on the second leaf of sorghum plants for monitoring their feeding behavior, as previously described (Cardona et al., 2022; Tetreault et al., 2019). Each recording session included control and treatment aphids run concurrently, and experiments were repeated across multiple sessions over 7 months. Treatment and control recordings were distributed across experimental sessions. Eight-hour recordings are commonly used in aphid EPG studies and are sufficient to capture pathway, salivation, xylem feeding, and phloem ingestion events. We used the EPG acquisition software *Stylet+* (EPG Systems, Wageningen, The Netherlands) to record SCA waveforms. Waveforms were annotated and quantified for all replicates and categorized according to standard EPG waveform definitions as pathway, non-probing, E1 (salivation), E2 (phloem ingestion), E1-E2 transitions, xylem, FPR (time to first probe) and FPHL (time to first phloem phase) (Cardona et al., 2022; Tetreault et al., 2019). Waveform durations were recorded in seconds and converted to minutes or hours where appropriate (Figures 2 and 3). Waveform durations were analyzed separately for each treatment relative to its corresponding control using Mann-Whitney tests for non-parametric data. Statistical significance was determined at *P* < 0.05 using GraphPad Prism version 10 (GraphPad Software, San Diego, CA, USA).

## Results

### ATPase assay

ATPase assays showed that both treatments significantly decreased the ATPase activity relative to aphids fed on the control diet (Figure 1; F-value = 7.726, *P* = 0.0020). Related to the control, ATPase activity was reduced in dexamethasone-tested aphids *(Padj* dexamethasone = 0.0188; mean difference = 0.001740) and elacridar-treated aphids (*Padj* elacridar = 0.0024; with mean differences = 0.002405). ATPase activities did not differ significantly between dexamethasone- and elacridar-treated groups (*Padj =* 0.5686; mean difference = 0.0006650).

**Figure 1:**
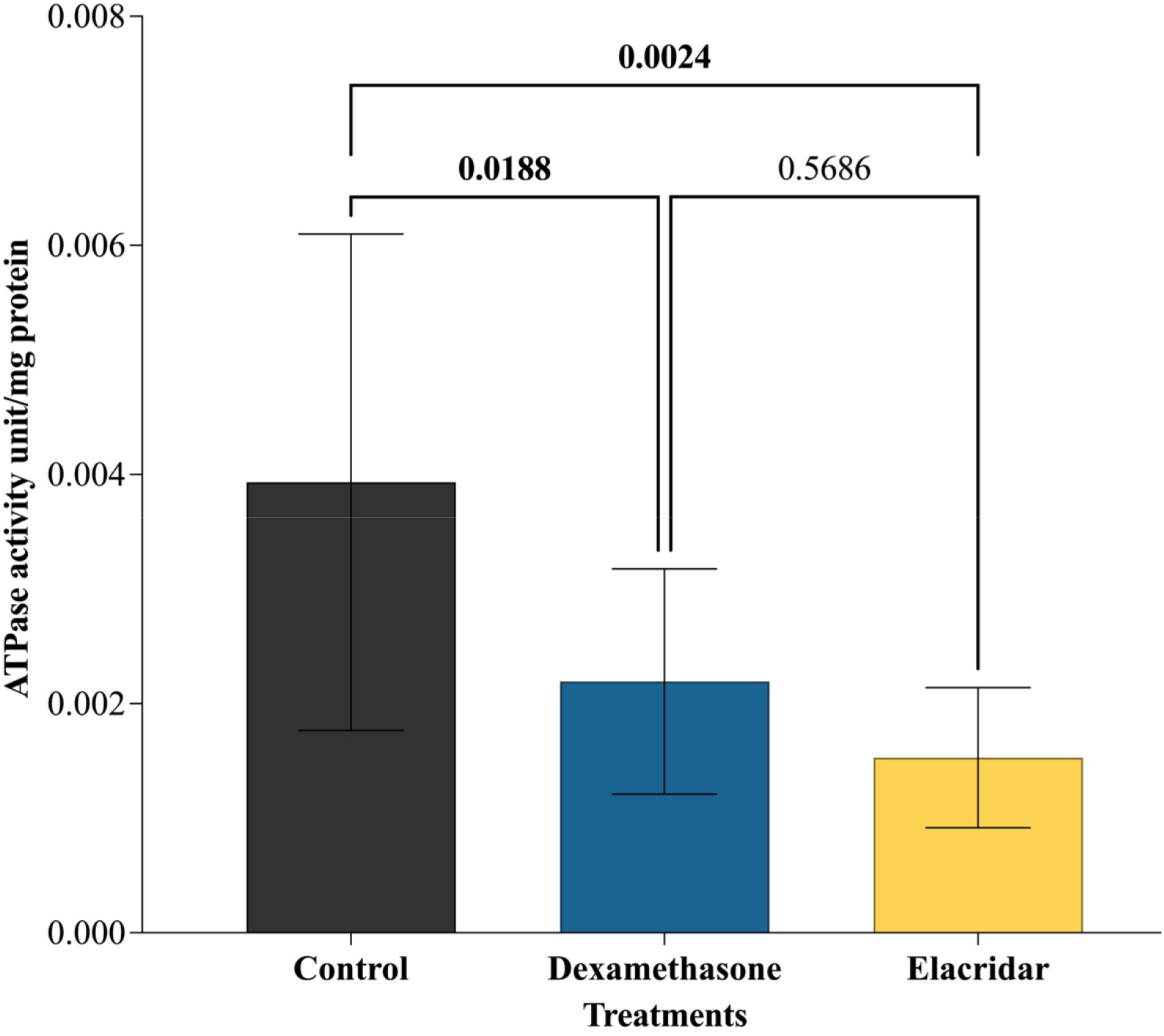
ATPase activity after 24 hours on treated diets, measured at 620 nm, comparison between the control diet, dexamethasone-supplemented diet and elacridar-supplemented diet. Mean ATPase activity +/-SD, *P*-values from one-way ANOVA and Tukey’s multiple comparisons test.

### Electrical Penetration Graph

To monitor the effect of ABC transporter modulators on SCA feeding behavior, we collected data for 13 replicates of dexamethasone-treated aphids, along with their controls (n = 10), and 12 replicates of elacridar-treated aphids, along their controls (n = 11). While several comparisons were not significantly different between controls and treatments (0.9415 > *P* > 0.1808; Table 1, Supplemental Figures S2 and S3), three comparisons showed significant variations between dexamethasone- and elacridar-treated SCA compared to respective controls. For dexamethasone-fed SCA, the pathway phase duration was significantly increased compared to the untreated individuals (*P* = 0.0422; Sum of ranks in control vs dexamethasone: 87 vs 189; Figure 2. Table 1). For elacridar-treated SCA, the salivation E1 phase was significantly decreased compared to the untreated individuals (*P*= 0.0225; Sum of ranks in control vs elacridar: 169 vs 107, Figure 3, Table 1). Moreover, the xylem phase duration was significantly increased for elacridar-treated SCA compared to the untreated individuals (*P*= 0.0012; Sum of ranks in control vs elacridar: 85.50 vs 190.5) as shown in Figure 3 and Table 1. Representative waveforms for each affected phase in controls and treated SCA are represented in Suplemental Figure S4.

**Figure 2:**
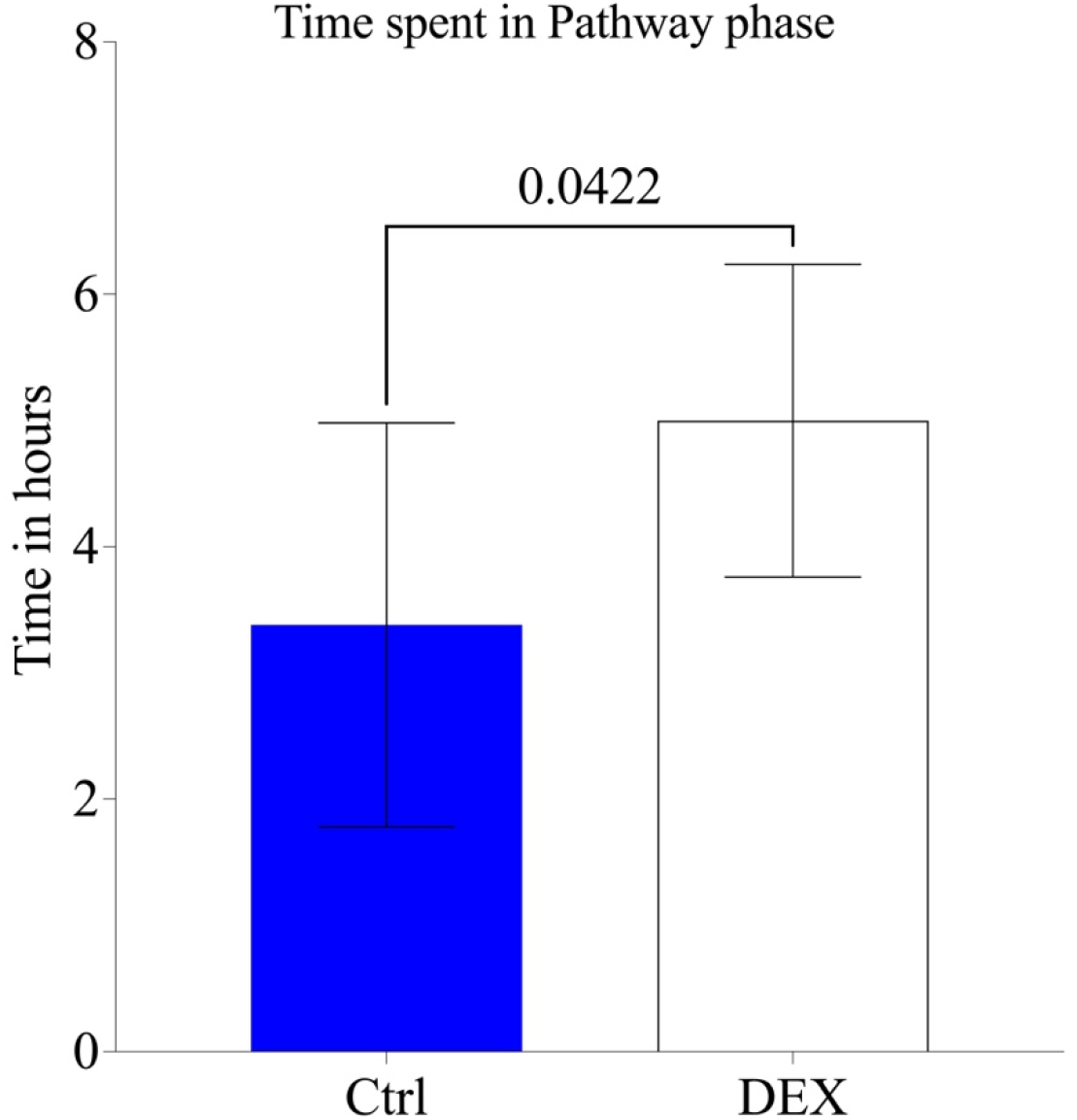
Comparison of the time spent, in hours, in pathway phase for aphids fed on control diet (Ctrl: sucrose) and dexamethxasone diet (sucrose-dexamethasone 0.5 mM, DEX). The difference in total duration was significant, mean time +/-SD, *P* = 0.0422, Mann-Whitney tests for non-parametric data.

**Figure 3:**
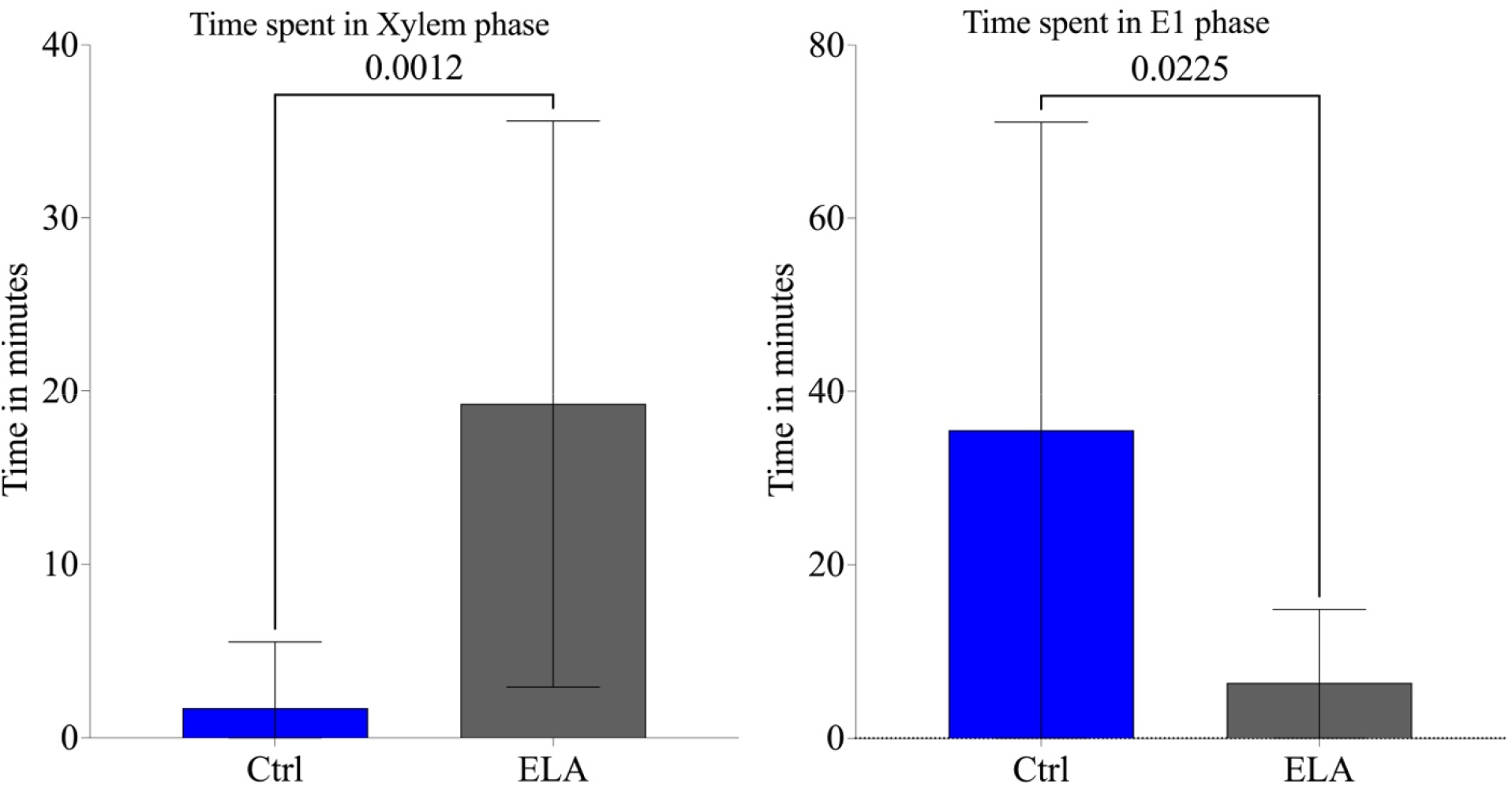
Comparison of the time spent, in minutes, in xylem phase (left) and E1 phase (right) for aphids fed on control diet (Ctrl: sucrose) and elacridar diet (sucrose-elacridar 0.5 mM, ELA), with mean time +/-SD, Mann-Whitney tests for non-parametric data.

**Table 1:** Effects of ABC transporter modulators on sugarcane aphid EPG feeding phases. Values are medians. *P*-values are from two-tailed Mann–Whitney tests comparing each treatment with its corresponding control.

| Feeding parameter | Control median | Treatment median | n Control | n Treatment | U | *P*-value | Significance |
| --- | --- | --- | --- | --- | --- | --- | --- |
| **Dexamethasone** |  |  |  |  |  |  |  |
| No probing | 13.1 min | 14.85 min | 10 | 13 | 63.5 | 0.9415 | Ns |
| Pathway | 3.686 h | 5.132 h | 10 | 13 | 32.0 | 0.0422 | * |
| E1 salivation | 7.39 min | 18.12 min | 10 | 13 | 48.0 | 0.3128 | Ns |
| E1–E2 transition | 6.23 min | 13.52 min | 10 | 13 | 43.0 | 0.1808 | Ns |
| E2 phloem ingestion | 3.57 h | 1.63 h | 10 | 13 | 47.5 | 0.2909 | Ns |
| Xylem | 3.29 min | 0.0 s | 10 | 13 | 52.5 | 0.4075 | Ns |
| FPR | 1.0 s | 1.0 s | 10 | 13 | 53.5 | 0.3950 | Ns |
| FPHL | 1.43 h | 2.09 h | 10 | 13 | 49.0 | 0.3434 | Ns |
| **Elacridar** |  |  |  |  |  |  |  |
| No probing | 16.5 min | 40.32 min | 11 | 12 | 51.0 | 0.3793 | Ns |
| Pathway | 4.98 h | 4.64 h | 11 | 12 | 59.0 | 0.6947 | Ns |
| E1 salivation | 27.03 min | 4.62 min | 11 | 12 | 29.0 | 0.0225 | * |
| E1–E2 transition | 12.85 min | 5.28 min | 11 | 12 | 45.0 | 0.2115 | ns |
| E2 phloem ingestion | 1.48 h | 19.45 min | 11 | 12 | 49.0 | 0.3164 | ns |
| Xylem | 0.0 min | 16.01 min | 11 | 12 | 19.5 | 0.0012 | ** |
| FPR | 1.0 s | 1.0 s | 11 | 12 | 61.5 | 0.8238 | ns |
| FPHL | 1.496 h | 2.15 h | 11 | 12 | 63.0 | 0.8801 | ns |

ns, not significant; \**P* < 0.05; \*\**P* < 0.01. FPR, time to first probe; FPHL, time to first phloem. Sources: dexamethasone and elacridar Mann–Whitney outputs.

## Discussion

This study used the sugarcane aphid (SCA) as a model for piercing-sucking herbivory to examine whether ABC transporter modulators can influence aphid feeding on sorghum. We hypothesized that pretreatment with these modulators would alter SCA feeding behavior possibly by changing ABC transporter function and activity. Accordingly, modulator-induced changes in feeding would provide indirect evidence that ABC transporters contribute to the feeding process. Our results show that ABC transporter modulators affect specific phases of SCA feeding rather than causing a general cessation of feeding. This pattern suggests that altered transporter activity selectively disrupts distinct components of feeding behavior, supporting a mechanistic role of ABC transporters in coordinating aphid feeding on sorghum. We further showed reduced ATPase activity in aphids exposed to either transporter modulator relative to untreated aphids. This result was expected for elacridar, a well-established inhibitor of ABC transporter activity that has been shown to reduce transporter-associated ATPase activity in human cell lines (Hamaguchi-Suzuki et al., 2024; Stasiak et al., 2025). In contrast, dexamethasone is commonly characterized as an inducer or activator of ABC transporter expression and activity in mammalian systems (Narang et al., 2008; Ueda et al., 1992). However, the reduced ATPase activity observed in dexamethasone-exposed aphids suggests that its effects on insect ABC transporters may differ from those reported in vertebrates. These differences may reflect taxon-specific variation in transporter structure, regulation, or tissue-specific expression. Alternatively, dexamethasone may interact with different ABC transporter subfamilies in insects than those characterized in mammalian systems. Together, these findings highlight the need to identify the specific transporter families and physiological pathways affected by dexamethasone in aphids.

The two modulators produced distinct effects on SCA feeding behavior. Dexamethasone increased the duration of the pathway phase, whereas elacridar reduced E1 salivation and increased xylem feeding duration. Because the E1 phase, which corresponds to watery saliva secretion (Grover et al., 2020), is associated with the release of a non-persistent virus in plant tissues (Martín et al., 1997; Powell, 2005), reducing the time aphids spend salivating may be particularly relevant for managing aphid-transmitted viruses. These distinct behavioral phenotypes likely reflect different physiological consequences of ABC transporter modulation. Because dexamethasone and elacridar may affect different ABC transporter systems, individual feeding phases may depend on distinct transporter-mediated processes, including ion homeostasis, metabolite trafficking, salivary secretion, and mitigation of cellular stress. The increased xylem feeding observed following elacridar exposure, particularly in combination with reduced E1 salivation, may indicate disruption of salivary gland function or feeding-related osmoregulation. ABC transporters could support salivary gland physiology by regulating ion gradients, transporting metabolites, lipids, or peptides, and maintaining cellular homeostasis during sustained feeding. Aphid salivation requires tightly coordinated secretion during stylet penetration and phloem contact (Tjallingii, 2006); therefore, inhibition of transporter activity could alter salivary composition or secretion efficiency, disrupting normal feeding progression. Alternatively, increased xylem feeding may reflect osmotic stress or difficulty maintaining phloem ingestion. A similar response has been reported in the Asian citrus psyllid, *Diaphorina citri* (Kuwayama), in which increased xylem ingestion was associated with impaired phloem feeding on resistant host plants (Shugart et al., 2019). Additionally, SCA spent more time feeding on the *bloomless* mutant (lacking waxes) than on wild-type sorghum plants, a response associated with water stress, counteracted by consuming more water from plant xylem to maintain water balance in the gut (Cardona et al., 2022).

In contrast, dexamethasone increased the time SCA spent in pathway phase. This phase encompasses stylet penetration and intercellular navigation as aphids search for a suitable ingestion site while forming a salivary sheath (Backus et al., 2020). This prolonged pathway phase may therefore reflect extended searching for accessible phloem cells, impaired salivary sheath formation, altered navigation through host tissues, or delayed progression to sustained phloem feeding. Similar behaviors have been observed when hemipteran insects encounter plant traits that impede feeding. For example, in *D. citri*, for instance, prolonged pathway activities have been associate with physical barriers, including a fibrous ring surrounding phloem cells (George et al., 2017; Shugart et al., 2019). Extended probing also occurs when aphids feed on non-host plants or encounter other plant characteristics that reduce feeding success (Escudero-Martinez et al., 2021). Notably, the increased pathway duration observed here cannot be attributed to plant resistance, because aphids fed on the same sorghum host plants across treatments. Instead, dexamethasone exposure appears to have altered aphid physiology in a manner that impaired or delayed progression through the pathway phase.

The contrasting effects of dexamethasone and elacridar suggest that different ABC transporter families, or distinct transporter-associated physiological pathways, contribute to specific aspects of aphid feeding behavior. The divergent feeding phenotypes therefore support the hypothesis that aphid feeding is regulated by a balance of transporter activities rather than by a simple on-off mechanism.

Although significant alterations were observed in the pathway, salivation, and xylem-feeding phases, sustained phloem ingestion was comparatively less affected under our experimental conditions. This pattern suggests that ABC transporter modulation may primarily disrupt early feeding coordination, host-tissue navigation, or physiological compensation rather than phloem ingestion itself. Alternatively, stronger effects on E2 phloem-ingestion phase may require longer exposure periods or more extensive transporter inhibition.

Together, these findings indicate that ABC transporter activity contributes to multiple physiological processes underlying SCA feeding behavior. Although ABC transporters have traditionally been studied in the context of insecticide detoxification and resistance, our results support a broader role for these proteins in hemipteran feeding physiology. Multifunctional roles of ABC transporters have also been reported in other insects, including nutrient transport, stress responses, epithelial homeostasis, and interactions with host-derived compounds (Bretschneider et al., 2016; Enders et al., 2020; Hull et al., 2014; Jin et al., 2020; Liu et al., 2019; Wang et al., 2023).

Pharmacological modulation provides a useful initial approach to examining transporter function; however, these compounds may affect multiple ABC transporter subfamilies and potentially influence additional physiological pathways. Future work should therefore identify the specific transporters involved through transcriptomic analyses, tissue-specific expression profiling, RNA interference, or gene editing approaches. Direct measurements of virus transmission following transporter modulation will also be important for determining whether reduced E1 salivation duration translates into lower transmission efficiency under biologically relevant conditions. Collectively, our findings demonstrate that ABC transporter modulators do disrupt multiple feeding phases associated with plant damage and pathogen transmission of aphids. Targeting physiological regulators of feeding behavior may offer an alternative to strategies that rely exclusively on lethality. Such approaches could reduce crop injury and limit virus spread while potentially lowering the selection pressure associated with conventional insecticides.

## References

Backus, E.A., Guedes, R.N.C., Reif, K.E., 2020. AC–DC electropenetrography: fundamentals, controversies, and perspectives for arthropod pest management. 10.1002/ps.6087

Bansal, R., Mian, M., Mittapalli, O., Michel, A., 2014. RNA-Seq reveals a xenobiotic stress response in the soybean aphid, Aphis glycines, when fed aphid-resistant soybean. BMC Genomics 15, 972.

Bhargava, K.S., Joshi, R.D., Rizvi, S.M.A., 1971. Some observations on the insect transmission of sugarcane mosaic virus. Sugarcane Pathol Newslett.

Bowling, R.D., Brewer, M.J., Kerns, D.L., Gordy, J., Seiter, N., Elliott, N.E., Buntin, G.D., Way, M.O., Royer, T.A., Biles, S., Maxson, E., 2016. Sugarcane aphid (Hemiptera: Aphididae): A new pest on sorghum in North America. Journal of Integrated Pest Management 7, 12. 10.1093/jipm/pmw011

Brault, V., Uzest, M., Monsion, B., Jacquot, E., Blanc, S., 2010. Aphids as transport devices for plant viruses. Comptes Rendus Biologies, Les puceron□: modèles biologiques et ravageurs des cultures 333, 524–538. 10.1016/j.crvi.2010.04.001

Bretschneider, A., Heckel, D.G., Vogel, H., 2016. Know your ABCs: Characterization and gene expression dynamics of ABC transporters in the polyphagous herbivore Helicoverpa armigera. Insect Biochemistry and Molecular Biology 72, 1–9. 10.1016/j.ibmb.2016.03.001

Cardona, J.B., Grover, S., Busta, L., Sattler, S.E., Louis, J., 2022. Sorghum cuticular waxes influence host plant selection by aphids. Planta 257, 22. 10.1007/s00425-022-04046-3

Dahlberg, J., 2019. The role of sorghum in renewables and biofuels, in: Zhao, Z.-Y., Dahlberg, J. (Eds.), Sorghum: Methods and Protocols, Methods in Molecular Biology. Springer, New York, NY, pp. 269–277.

Dermauw, W., Van Leeuwen, T., 2014. The ABC gene family in arthropods: Comparative genomics and role in insecticide transport and resistance. Insect Biochemistry and Molecular Biology 45, 89–110. 10.1016/j.ibmb.2013.11.001

EDDMapS, 2026. Sugarcane aphid (Melanaphis sacchari (Zehntner, 1897)) - EDDMapS State Distribution [WWW Document]. EDDMapS.org. URL https://www.eddmaps.org/distribution/usstate.cfm?sub=8170 (accessed 9.4.26).

Enders, L.S., Rault, L.C., Heng-Moss, T.M., Siegfried, B.D., Miller, N.J., 2020. Transcriptional responses of soybean aphids to sublethal insecticide exposure. Insect Biochemistry and Molecular Biology 118, 103285. 10.1016/j.ibmb.2019.103285

Escudero-Martinez, C., Leybourne, D.J., Bos, J.I.B., 2021. Plant resistance in different cell layers affects aphid probing and feeding behaviour during non-host and poor-host interactions. Bulletin of Entomological Research 111, 31–38. 10.1017/S0007485320000231

Gampa, G., Talele, S., Kim, M., Mohammad, A., Griffith, J., Elmquist, W.F., 2020. Chapter 9 - Influence of transporters in treating cancers in the CNS, in: Sosnik, A., Bendayan, R. (Eds.), Drug Efflux Pumps in Cancer Resistance Pathways: From Molecular Recognition and Characterization to Possible Inhibition Strategies in Chemotherapy, Cancer Sensitizing Agents for Chemotherapy. Academic Press, pp. 277–301. 10.1016/B978-0-12-816434-1.00009-7

George, J., Ammar, E.-D., Hall, D.G., Lapointe, S.L., 2017. Sclerenchymatous ring as a barrier to phloem feeding by Asian citrus psyllid: Evidence from electrical penetration graph and visualization of stylet pathways. PLOS ONE 12, e0173520. 10.1371/journal.pone.0173520

Grover, S., Varsani, S., Kolomiets, M.V., Louis, J., 2020. Maize defense elicitor, 12-oxo-phytodienoic acid, prolongs aphid salivation. Communicative & Integrative Biology 13, 63–66. 10.1080/19420889.2020.1763562

Hamaguchi-Suzuki, N., Adachi, N., Moriya, T., Yasuda, S., Kawasaki, M., Suzuki, K., Ogasawara, S., Anzai, N., Senda, T., Murata, T., 2024. Cryo-EM structure of P-glycoprotein bound to triple elacridar inhibitor molecules. Biochemical and Biophysical Research Communications 709, 149855. 10.1016/j.bbrc.2024.149855

Heckel, D.G., Gahan, L.J., Baxter, S.W., Zhao, J.Z., Shelton, A.M., Gould, F., Tabashnik, B.E., 2007. The diversity of Bt resistance genes in species of Lepidoptera. Journal of Invertebrate Pathology 95, 192–197. 10.1016/j.jip.2007.03.008

Hull, J.J., Chaney, K., Geib, S.M., Fabrick, J.A., Brent, C.S., Walsh, D., Lavine, L.C., 2014. Transcriptome-based identification of ABC transporters in the Western tarnished plant bug Lygus hesperus. PLoS One 9. 10.1371/journal.pone.0113046

Jin, M., Cheng, Y., Guo, X., Li, M., Chakrabarty, S., Liu, K., Wu, K., Xiao, Y., 2020. Down-regulation of lysosomal protein ABCB6 increases gossypol susceptibility in Helicoverpa armigera. Insect Biochemistry and Molecular Biology 122, 103387. 10.1016/j.ibmb.2020.103387

Khalid, W., Ali, A., Arshad, M.S., Afzal, F., Akram, R., Siddeeg, A., Kousar, S., Rahim, M.A., Aziz, A., Maqbool, Z., Saeed, A., 2022. Nutrients and bioactive compounds of Sorghum bicolor L. used to prepare functional foods: A review on the efficacy against different chronic disorders. International Journal of Food Properties 25, 1045–1062. 10.1080/10942912.2022.2071293

Li, Z., Davis, J.A., Swale, D.R., 2019. Chemical inhibition of Kir channels reduces salivary secretions and phloem feeding of the cotton aphid, Aphis gossypii (Glover). Pest Management Science 75, 2725–2734. 10.1002/ps.5382

Liu, Y., Qi, M., Dietrich, C.H., He, Z., Wei, C., 2019. Comparative sialotranscriptome analysis of the rare Chinese cicada Subpsaltria yangi, with identification of candidate genes related to host-plant adaptation. International Journal of Biological Macromolecules 130, 323–332. 10.1016/j.ijbiomac.2019.02.132

Long, W.H., Hensley, S.D., 1972. Insect pests of sugar cane. Annual Review of Entomology 17, 149–176. 10.1146/annurev.en.17.010172.001053

Martín, B., Collar, J.L., Tjallingii, W.F., Fereres, A., 1997. Intracellular ingestion and salivation by aphids may cause the acquisition and inoculation of non-persistently transmitted plant viruses. Journal of General Virology 78, 2701–2705. 10.1099/0022-1317-78-10-2701

Narang, V.S., Fraga, C., Kumar, N., Shen, J., Throm, S., Stewart, C.F., Waters, C.M., 2008. Dexamethasone increases expression and activity of multidrug resistance transporters at the rat blood-brain barrier. Am J Physiol Cell Physiol 295, C440–C450. 10.1152/ajpcell.00491.2007

O’Hara, F.M., Liu, Z., Davis, J.A., Swale, D.R., 2023. Catalyzing systemic movement of inward rectifier potassium channel inhibitors for antifeedant activity against the cotton aphid, Aphis gossypii (Glover). Pest Management Science 79, 194–205. 10.1002/ps.7188

Powell, G., 2005. Intracellular salivation is the aphid activity associated with inoculation of non-persistently transmitted viruses. Journal of General Virology 86, 469–472. 10.1099/vir.0.80632-0

Schenck, S., Lehrer, A.T., 2000. Factors affecting the transmission and spread of sugarcane yellow leaf virus. Plant Disease 84, 1085–1088. 10.1094/PDIS.2000.84.10.1085

Shugart, H., Ebert, T., Gmitter, F., Rogers, M., 2019. The Power of Electropenetrography in Enhancing Our Understanding of Host Plant-Vector Interactions. Insects 10. 10.3390/insects10110407

Singh, B.U., Padmaja, P.G., Seetharama, N., 2004. Biology and management of the sugarcane aphid, Melanaphis sacchari (Zehntner) (Homoptera: Aphididae), in sorghum: A review. Crop Protection 23, 739–755. 10.1016/j.cropro.2004.01.004

Smith, P.K., Krohn, R.I., Hermanson, G.T., Mallia, A.K., Gartner, F.H., Provenzano, M.D., Fujimoto, E.K., Goeke, N.M., Olson, B.J., Klenk, D.C., 1985. Measurement of protein using bicinchoninic acid. Analytical Biochemistry 150, 76–85. 10.1016/0003-2697(85)90442-7

Stasiak, P., Sopel, J., Płóciennik, A., Musielak, O., Lipowicz, J.M., Rawłuszko-Wieczorek, A.A., Sterzyńska, K., Korbecki, J., Januchowski, R., 2025. Elacridar Inhibits BCRP Protein Activity in 2D and 3D Cell Culture Models of Ovarian Cancer and Re-Sensitizes Cells to Cytotoxic Drugs. Int J Mol Sci 26, 5800. 10.3390/ijms26125800

Tarr, P.T., Tarling, E.J., Bojanic, D.D., Edwards, P.A., Baldán, Á., 2009. Emerging new paradigms for ABCG transporters. Biochimica et Biophysica Acta (BBA) - Molecular and Cell Biology of Lipids, Cellular Lipid Transport Processes and their Role in Human Disease 1791, 584–593. 10.1016/j.bbalip.2009.01.007

Tetreault, H.M., Grover, S., Scully, E.D., Gries, T., Palmer, N.A., Sarath, G., Louis, J., Sattler, S.E., 2019. Global responses of resistant and susceptible sorghum (Sorghum bicolor) to sugarcane aphid (Melanaphis sacchari). Frontiers in Plant Science 10, 145. 10.3389/fpls.2019.00145

Tjallingii, W.F., 2006. Salivary secretions by aphids interacting with proteins of phloem wound responses. Journal of Experimental Botany 57, 739–745. 10.1093/jxb/erj088

Ueda, K., Okamura, N., Hirai, M., Tanigawara, Y., Saeki, T., Kioka, N., Komano, T., Hori, R., 1992. Human P-glycoprotein transports cortisol, aldosterone, and dexamethasone, but not progesterone. J. Biol. Chem. 267, 24248–24252.

Vasquez, A., Belsky, J., Khanal, N., Puri, H., Balakrishnan, D., Joshi, N.K., Louis, J., Studebaker, G., Kariyat, R., 2025. Melanaphis sacchari/sorghi complex: current status, challenges and integrated strategies for managing the invasive sap-feeding insect pest of sorghum. Pest Management Science 81, 2427–2441. 10.1002/ps.8291

Wang, Y.-H., Klobasa, W., Chu, F.-C., Huot, O., Whitfield, A.E., Lorenzen, M., 2023. Structural and functional insights into the ATP-binding cassette transporter family in the corn planthopper, Peregrinus maidis. Insect Molecular Biology 32, 412–423. 10.1111/imb.12840

Wille, B.D., Hartman, G.L., 2008. Evaluation of artificial diets for rearing Aphis glycines (Hemiptera: Aphididae). Journal of Economic Entomology 101, 1228–1232. 10.1093/jee/101.4.1228

Wright, R., Peterson, J., Bradshaw, J., 2016. Be on the lookout for sugarcane aphids on sorghum [WWW Document]. CropWatch. URL https://cropwatch.unl.edu/2016/be-lookout-sugarcane-aphids-sorghum (accessed 8.9.22).

