## Supplemental Figures for "Pharmacological modulation of ATP-binding cassette transporters alters feeding phases of the sugarcane aphid on susceptible sorghum"

* Sharing first authorship

### Supplemental Figure Captions

Supplemental Figure S1: SCA after diet feeding, with red coloring showing consumption of the treatments.

Supplemental Figure S2: Non-significant comparisons ($P > 0.05$) control vs dexamethasone EPG phases, represented as mean time +/- SD, from Mann-Whitney tests for non-parametric data.

Supplemental Figure S3: Non-significant comparisons ($P > 0.05$) control vs elacridar EPG phases, represented as mean time +/- SD, from Mann-Whitney tests for non-parametric data.

Supplemental Figure S4: Representative waveforms (volts) for the phases that differed between treatments over the 8-hour recordings, A. dexamethasone in top panel compared to its control, and B. elacridar with its control on the bottom panel. The phases that differed significantly after modulator consumption were highlighted, in blue for pathway and purple for xylem.

### Supplemental Figures

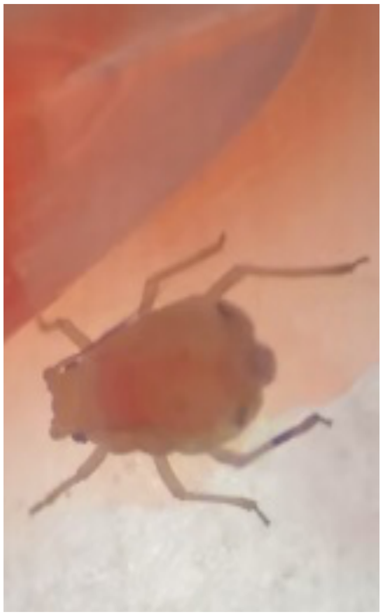

Supplemental Figure S1.

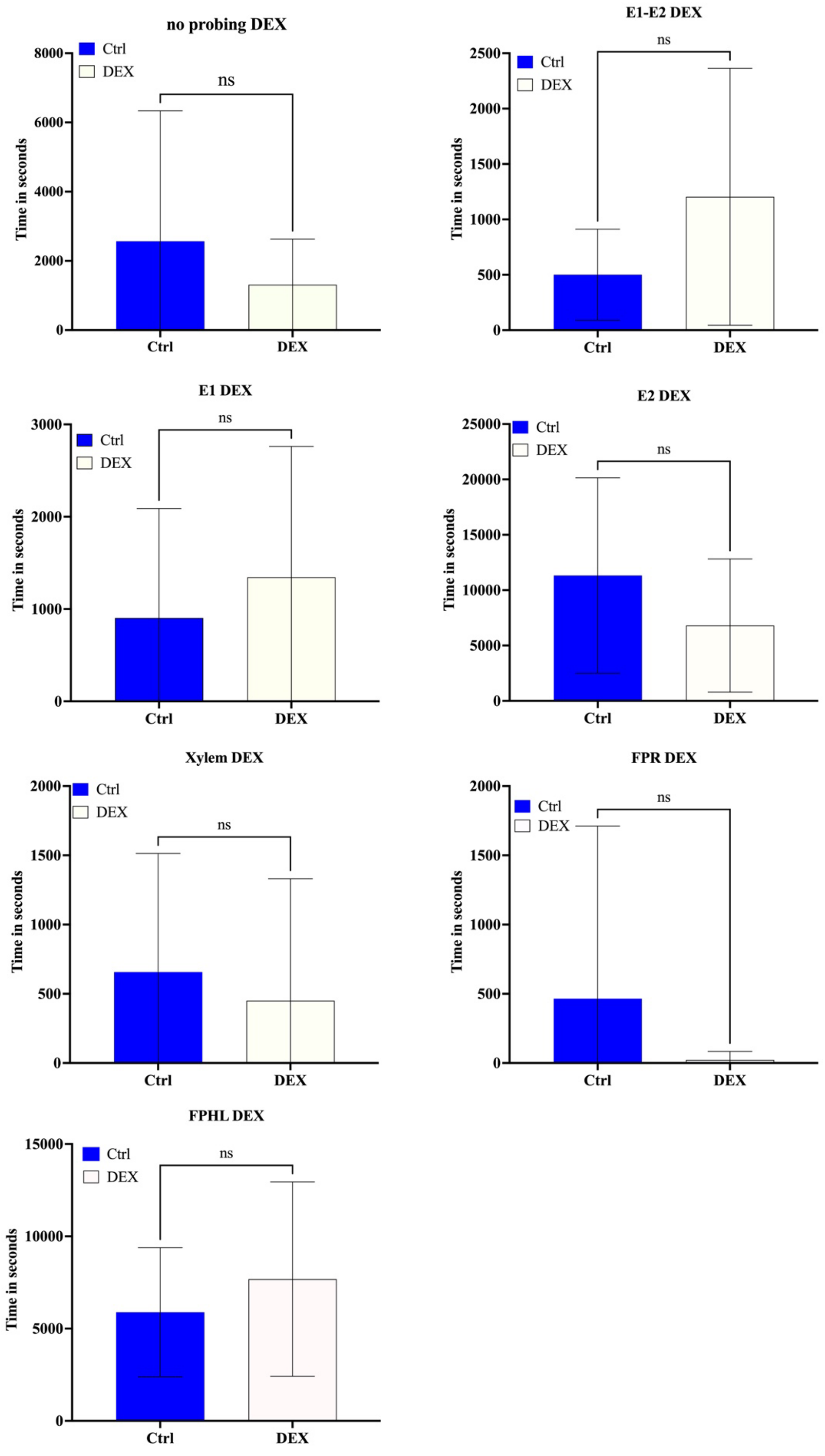


Supplemental Figure S2.

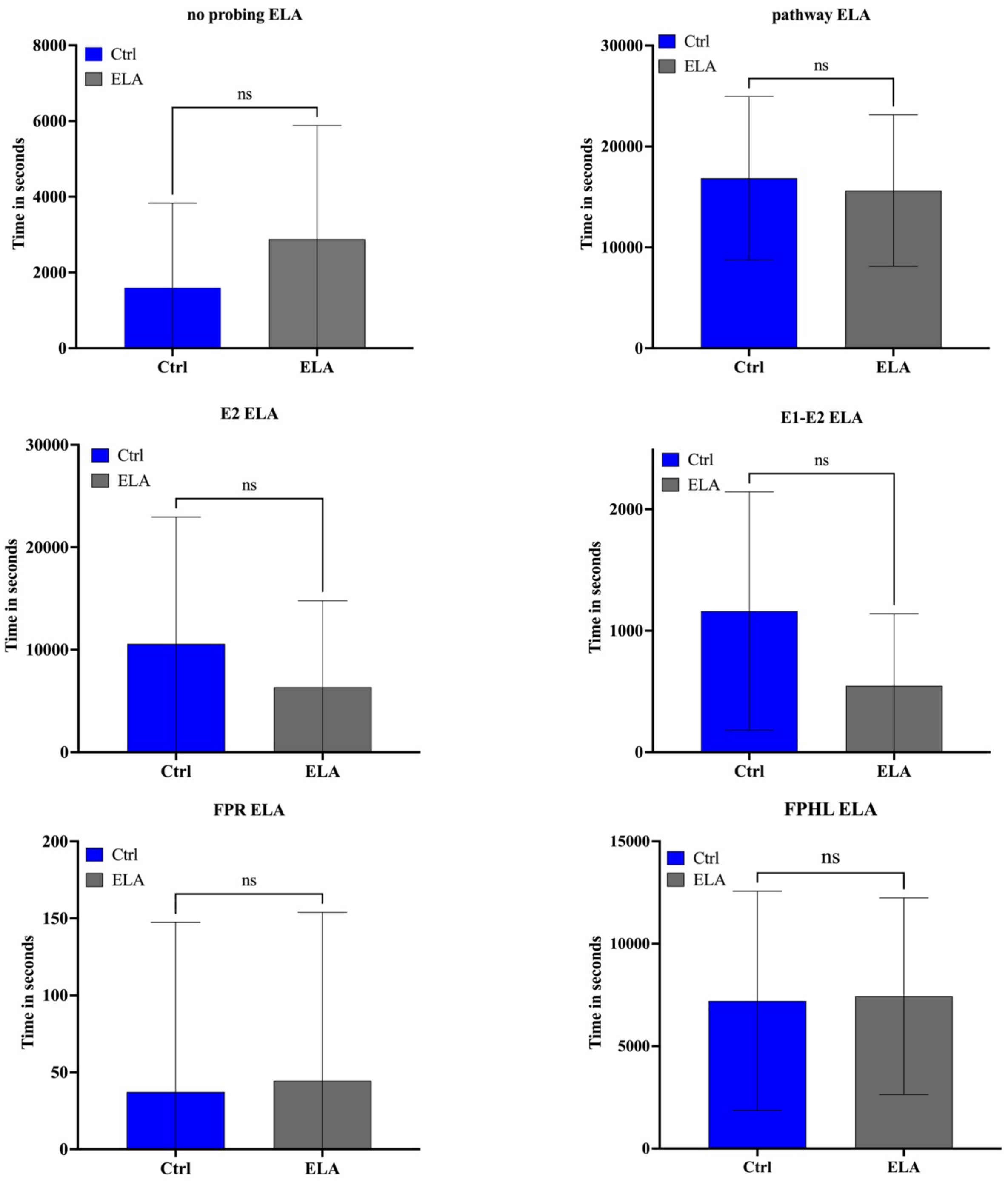


Supplemental Figure S3.

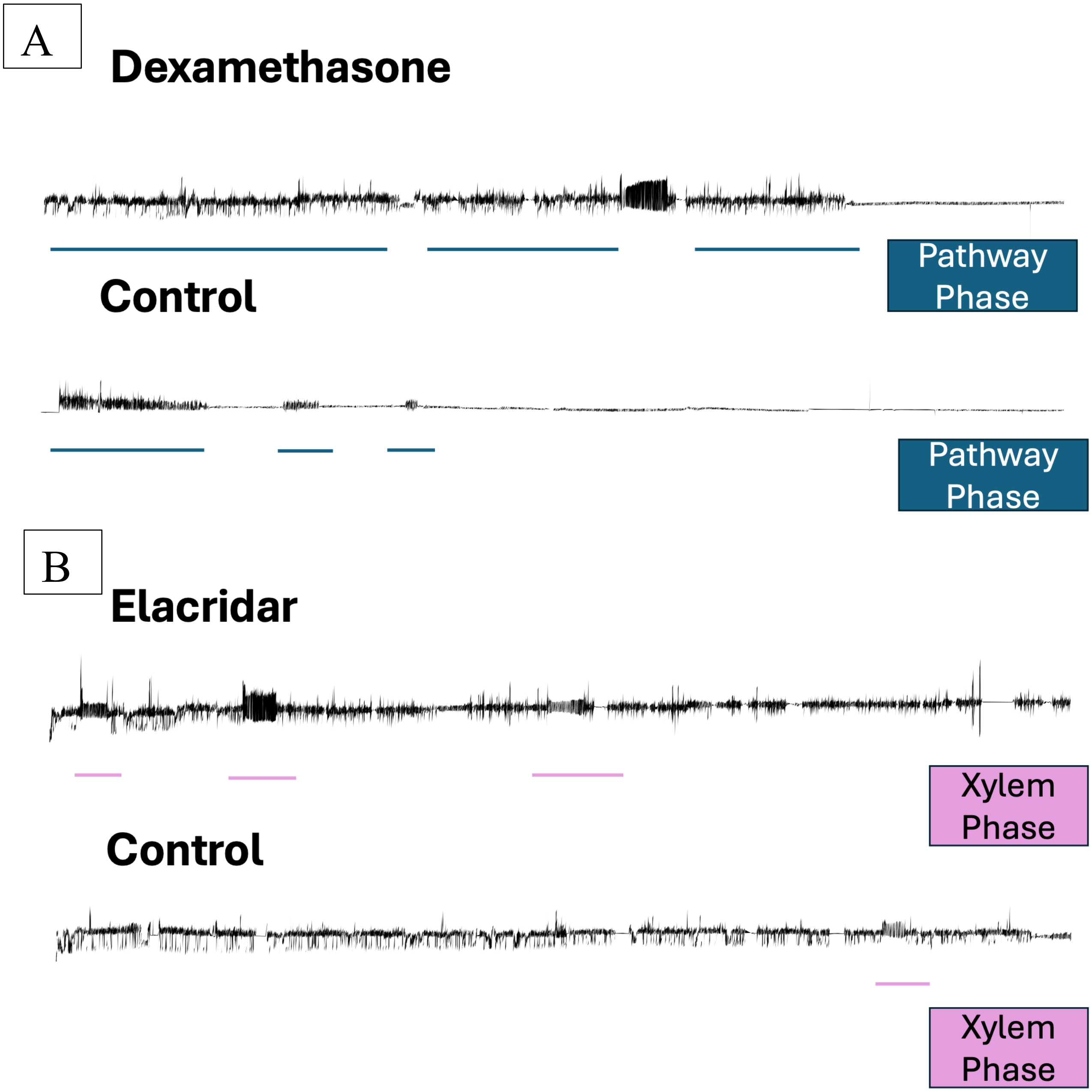


27 Supplemental Figure S4.
